# Killiverse: an interactive multi-omics web resource for killifish

**DOI:** 10.64898/2026.06.16.731504

**Authors:** Abhinav Mittal, Param Priya Singh

**Affiliations:** University of California, San Francisco, San Francisco, 94143, CA, USA; UCSF Bakar Aging Research Institute, San Francisco, CA, USA; UCSF Bakar Computational Health Science Institute, San Francisco, CA, USA

**Keywords:** killifish, multi-omics atlas, web application, aging, cross-study analysis

## Abstract

**Background:** Killifish have emerged as valuable vertebrate model systems for investigating several disciplines including aging, regeneration, and developmental biology. Multi-omics datasets are increasingly being generated for killifish. However, their reuse remains limited due to computational challenges, largely due to the lack of accessible resources creating a bottleneck in widespread adoption of the killifish model. To address this, we developed Killiverse, a web resource for quick and intuitive exploration of multi-modal omics data dedicated to the model organism.

**Results:** Killiverse is an interactive, no-code, web-based platform designed for exploration of killifish multi-omics data. The platform aggregates a growing list of datasets including bulk transcriptomes, single-cell and single-nucleus transcriptomes, proteomes, and lipidomes processed through standardized pipelines and genome assemblies. Killiverse supports customized visualization and enables cross-study and cross-species analysis. It provides ortholog mapping to several established model organisms. By combining low-code software development with modern cloud technologies, the platform delivers a scalable browser-accessible application for the community.

**Conclusions:** Killiverse enables rapid hypothesis development through the identification of patterns across studies and species. The ortholog maps allow the findings to be placed in a broader biological context. The platform represents an innovation in genomics data visualization that will serve as a template for future tool development. Killiverse is freely accessible at https://killiverse.org/.

## Background

Large-scale single-cell and multi-omics datasets generated across model organisms and humans have revolutionized our understanding of complex biological processes, but their reuse is often limited by fragmentation across repositories, heterogeneous processing pipelines, and the lack of interactive visualization tools. These challenges are particularly pronounced in emerging model systems such as killifish where datasets are growing rapidly but are scattered across data silos and remain difficult to explore in a unified and consistent manner. This limits the ability of researchers to fully leverage the unique biological properties of these model systems and to contextualize findings within the broader landscape of biomedical research.

Killifish are ray-finned fish in the order cyprinodontiformes comprising of more than 1200 species that can thrive in a wide variety of environments [1–3]. Their habitats range from the permanent rivers of South Asia, the alkaline desert pools of the southwestern United States to seasonal pools in Africa and South America. The killifish species that inhabit seasonal ponds (called annual killifish) have evolved unique adaptations including rapid sexual maturity, unique development trajectories and compressed lifespans making them popular models for developmental biology and aging research [2, 4]. For example, strains of the African turquoise killifish have remarkably short lifespan of 6-9 months making them a popular model for aging research. Even in this short lifespan they recapitulate many hallmarks of mammalian aging, including neurodegeneration, telomere attrition, loss of proteostasis, and chronic inflammation, making research findings directly relevant to human aging biology [5–8]. Killifish also possesses capacity for tissue regeneration and have been used to study regeneration of brain regions, optic nerves, tailfin and the loss of regenerative capacity with aging [9–13].

Annual killifish have a unique development due to their adaptation to the harsh seasonal habitat making them excellent models for stress resistance, and studies of ecological and evolutionary developmental biology. To survive the annual drought, their embryos can enter an extreme developmental dormancy called diapause at multiple developmental time points. Diapause embryos in the South American killifish *Austrofundulus limnaeus* can withstand environmental extreme including desiccation, oxidative stress and long periods of anoxia [14–16]. There is no tradeoff for future growth, fertility of lifespan even if the embryos stay in diapause for a period equal to adult killifish lifespan [17]. Therefore, understanding the molecular mechanisms of diapause can uncover new mechanisms of stress resistance, and longevity [18].

Due to these remarkable features, multiple species of killifish have emerged as valuable vertebrate model systems for investigating several disciplines [19–22], with a rapid accumulation of multimodal genomic datasets related to aging, diapause, regeneration and more. However, these data are scattered across individual studies, and their reuse and assessment of conservation of findings require substantial computational effort. To address this, we developed Killiverse, a freely accessible, web-based platform for exploring killifish multi-omics data. Killiverse hosts a growing collection of publicly available multi-modal omics datasets, processed through standardized pipelines and consistent genome assemblies. Detailed gene information and curated ortholog mappings to human, mouse, and other vertebrates further enable cross-species interpretation of findings. We aim to continue incorporating additional datasets and functionalities to establish Killiverse as a community resource for killifish genomics.

## Implementation

### Technical Architecture

Killiverse is developed using Dash (version 2.18.2), an open-source Python framework for building interactive web-based data applications (Figure 1). The platform is organized into modular dataset-specific pages, with each page implemented as a Python module defining layout components (e.g. dropdowns, graphs, and tables) and associated callback functions for interactivity. This study-specific design allows visualization types to be tailored to each dataset’s experimental structure while maintaining a consistent user interface across the application.

**Figure 1:**
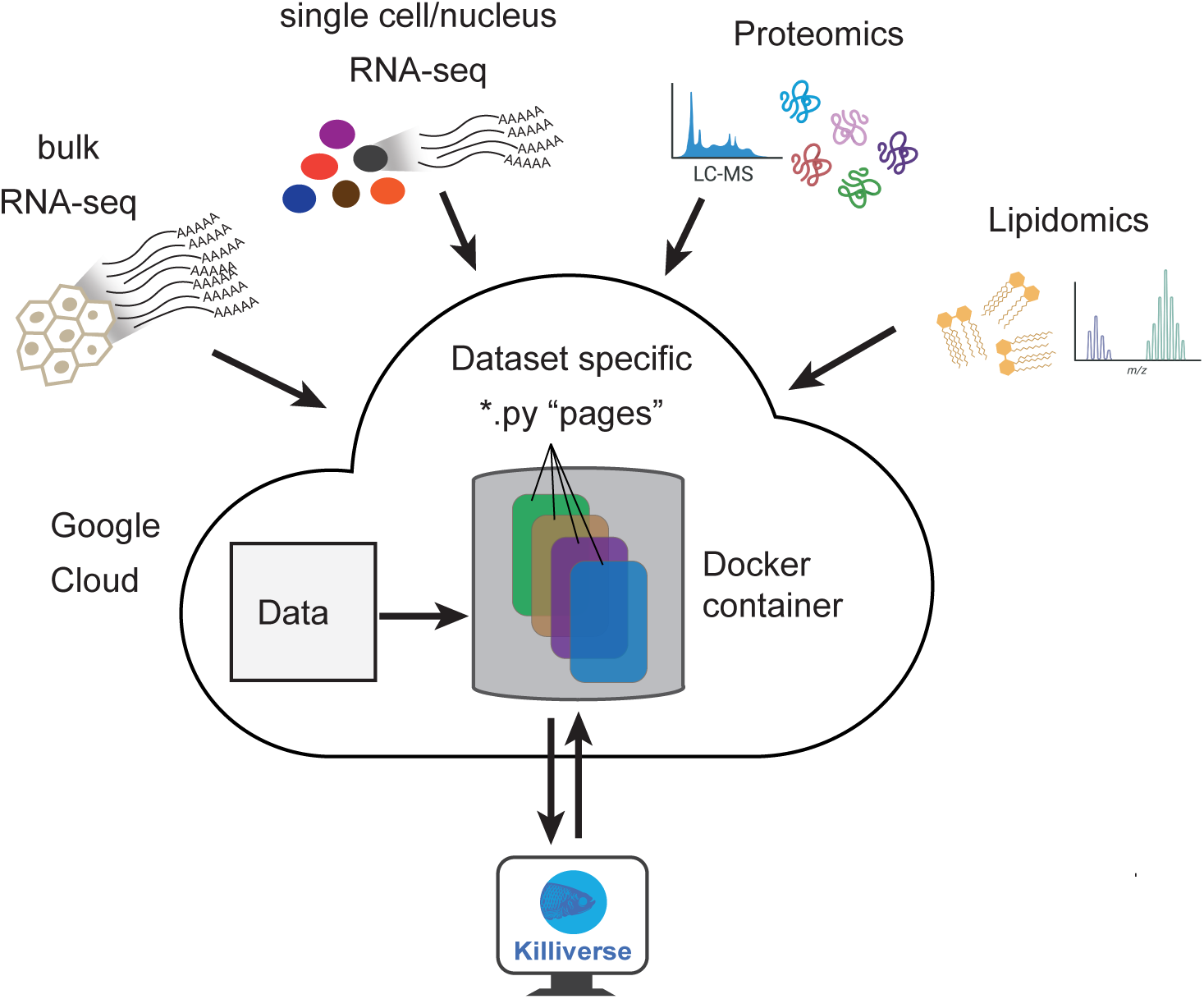
Technical architecture of Killiverse. Killiverse is developed using Dash framework in Python and deployed as a cloud application. Multi-modal omics data are processed and stored on google cloud in application specific python pages in a docker container. This architecture ensures custom study-specific visualization.

Data are stored either locally within the application or on Google Cloud Storage. The platform is deployed as a Docker container on Google Cloud Run, supporting reproducible deployment, scalability, and efficient resource management.

### Data acquisition and processing

Bulk RNA-seq datasets for killifish were obtained from NCBI GEO [23] and processed using the nf-core RNA-seq pipeline (version 3.18.0) implemented in Nextflow [24, 25]. Reads were aligned using STAR [26] and quantified using RSEM [27] against the respective genome assemblies [28, 29]. Study specific differential gene expression analysis was performed using PyDESeq2 [30]. This standardized processing ensures consistency and comparability across studies.

Single-cell and single-nucleus RNA-seq datasets were incorporated as processed AnnData [31] or Seurat [32] objects obtained from the original studies to preserve published annotations, dimensionality reductions, and cell type labels. Proteomics and lipidomics datasets were sourced as processed data from the original studies, including normalized abundance values and associated metadata.

### Navigating the application

The landing page provides brief introduction and outlines the significance of killifish as model organisms. Users can navigate to ‘Datasets (killifish)’ page that lists all the studies with brief description of the dataset. Studies and datasets can be filtered based on metadata such as assay type, species/strain, sex, and tissue (Figure S1). In addition to killifish datasets, Killiverse also hosts multi-tissue bulk RNA-seq datasets from mouse [33] and human [34] to support cross-species comparison, which can be accessed through ‘Datasets (human and mouse)’ tab in the navigation bar. The ‘Gene Information’ page provides information about orthologs in several species, and description of killifish orthologs in human and link to NCBI Gene pages in several species. ‘Help’ page has detailed instructions for the available functionalities and interactive features.

Across the platform, the tables support sorting, filtering, and rearrangement of columns, and export as CSV files. All the visualizations are interactive and downloadable as publication quality image in SVG format using icons on the top right of the visual elements.

### Dataset Exploration

Killiverse hosts a growing collection of publicly available datasets (Supplementary Table S1) spanning four modalities: bulk RNA-seq, single-cell and single-nucleus RNA-seq, lipidomics and proteomics, covering multiple killifish species and strains, tissues, and experimental designs including perturbation studies and aging time-courses. For comparison, one multi-tissue aging dataset each from mouse and human is also included, enabling direct cross-species inspection of age-associated expression changes.

Cross-study comparisons are confounded by differences in pipelines and genome assemblies across studies. Killiverse addresses this by uniformly reprocessing all bulk RNA-seq datasets through a single pipeline and genome assembly, enabling direct cross-study comparisons without significant computational effort.

For bulk RNA-seq studies, Killiverse provides differential gene expression analysis with distinct visualizations including volcano plots, box plots and heatmaps (Figure S2 and S3) and the ability to filter relevant comparisons based on p-value and log2 fold-change thresholds, and label genes of interest. Quantitative tables provide normalized counts, log2 fold-changes, and adjusted p-values for all genes, downloadable for downstream analysis. Heatmap tabs also allows to generate custom heatmaps of Z-scores from normalized counts for selected tissue/condition and any set of genes.

Eight of the bulk RNA-seq studies span four or more age groups (including mouse and human data), enabling longitudinal analysis of gene expression across the lifespan (Figure S5). For the longitudinal datasets, in addition to the heatmaps and boxplots, Killiverse also provides line graph views with a linear regression fit, reporting slope, R², and and p-value for each gene, allowing rapid triage of candidates with consistent age-associated expression changes (Figure S5). A ranked table of Spearman correlation coefficients (Figure S9) between expression and age allows to identify the strongest age-correlated genes across any tissue or condition.

For the single-cell and single-nucleus RNA-seq datasets, Killiverse provides views designed around the core challenges of cell-type-resolved expression analysis. These views allow researchers to assess whether a gene of interest is expressed ubiquitously or is restricted to a specific cell population, information critical for interpreting bulk RNA-seq results and designing follow-up experiments. The UMAP/t-SNE view renders each cell colored either by its annotation or by the expression level of a chosen gene, enabling context for both cluster identity and gene activity (Figure S13 and S14). The bubble plot view displays mean expression and the fraction of expressing cells for a user-defined gene set across all cell types simultaneously (Figure S15), and proportion plots summaries cell-type composition across samples or conditions, important in studies comparing tissues across ages or sexes where shifts in cellular composition can confound transcript-level comparisons. Quantitative tables report mean expression, positive cell fraction, and cell counts per annotation.

### Orthologs and Gene Information

To facilitate the identification of biologically relevant findings, particularly for killifish genes annotated with non-intuitive locus tags, Killiverse includes curated ortholog mappings to eight vertebrate species identified using reciprocal best-hit BLAST analysis [35] (see Supplementary Methods). For any selected gene, the platform also provides the human ortholog’s functional description from UniProt [36] and GO annotations from the PAN-GO Human Functionome [37]. Quality metrics for ortholog mapping, including query coverage, percent identity, and e-values, are also reported allowing users to assess mapping reliability before drawing functional inferences. This functionality facilitates cross-species interpretation of killifish findings and supports translation to human biology.

## Results

To demonstrate the utility of Killiverse for biological discovery, we explored how integration of datasets across species, strains, modalities, and life stages could uncover conserved molecular patterns that are not readily apparent from individual studies. The examples below illustrate how Killiverse can be used to identify shared regulatory programs across development, aging, and tissue homeostasis. The figures presented in the main text were generated by downloading data from Killiverse and replotting it for presentation purposes. The supplementary figures are semantically equivalent to the corresponding figures in the main text and were obtained directly from the platform as either downloaded SVG files or snapshots of the interactive visualizations, demonstrating the analyses and visualizations available within Killiverse.

### A conserved ezh1/ezh2 switch links diapause and aging across killifish species

To illustrate how Killiverse can enable easy cross-dataset discovery, we examined the expression of Polycomb Repressive Complex 2 (PRC2) members across killifish diapause and aging datasets. PRC proteins complexes are evolutionarily conserved regulators of chromatin state and gene silencing critical for development, stem cell identity, and cancer [38, 39]. PRC2 catalyzes the methylation of histone H3 on lysine 27 (H3K27me3), which is a hallmark of transcriptionally silent chromatin. Several members of PRC2 complex are highly upregulated during killifish diapause and are required for muscle maintenance during diapause in the African turquoise killifish *N. furzeri* [17]. The two catalytic subunits of PRC2 complex, ezh1 and ezh2, also have specialized expression during diapause and development respectively in the *N. furzeri* [40]. However, to what extant these findings are conserved across datasets, species and strains is unclear.

Using Killiverse, we identified five studies with bulk RNA-seq data during different stages of embryo development and diapause from two species, *N. furzeri* (GRZ and MZM-0403 strains) and *A. limnaeus* (Figure S1) [17, 28, 40, 41]. We found that in both the *N. furzeri* strains GRZ and MZM-0403, *ezh1* is consistently upregulated during diapause (Figures 2A, 2B, 2C and Figure S2) while *ezh2* shows elevated expression during pre-diapause and normal development (diapause escape or non-diapause) but not during diapause (Figures 2E, 2F, and 2G and Figure S2). This suggests a conserved *ezh2*-to-*ezh1* switch associated with diapause entry across different African turquoise killifish *N. furzeri* strains. RNA-seq data from development after embryos exit from diapause in the South American annual killifish *A. limnaeus* [42], show that the catalytic subunit switches back from *ezh1*-to-*ezh2* after embryos exit from diapause (Figures 2D and 2H).

**Figure 2.**
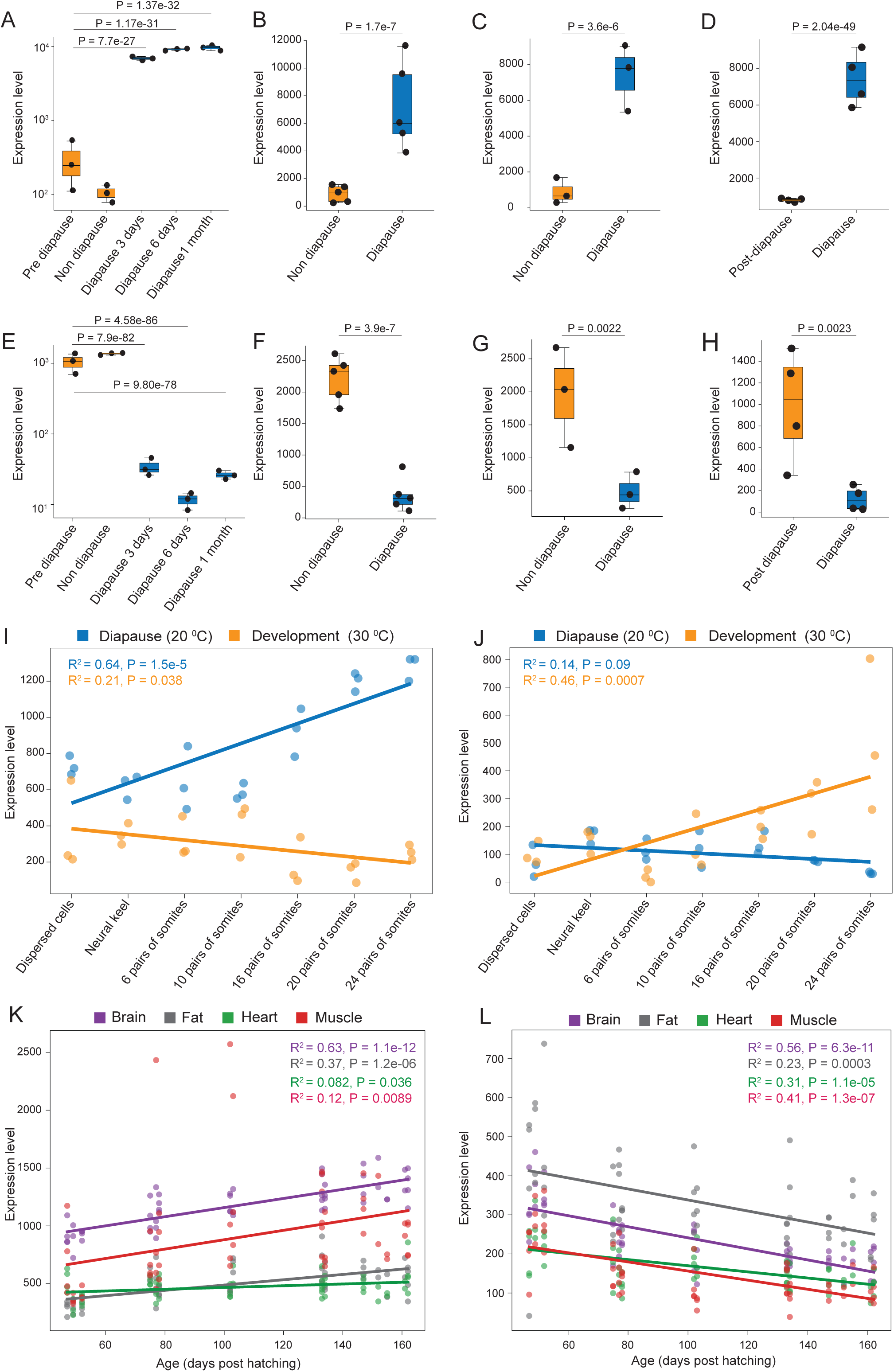
A conserved ezh1/ezh2 switch links diapause and aging across killifish species. Gene expression of the PRC2 complex catalytic subunits *ezh1* and *ezh2* from bulk RNA-seq datasets across killifish development, diapause and aging. **A.** *ezh1* expression in *N. furzeri* embryos (GRZ strain) during pre-diapause (onset of heartbeat), diapause escape (non-diapause development) and 3 days, 6 days and 1 month in diapause. **B, C.** *ezh1* expression during diapause and non-diapause development in *N. furzeri* (GRZ strain, B) and (MZM-0403 strain, C) from an independent study. **D.** *ezh1* expression during diapause and post-diapause development (4 days after exit from diapause) in *A. limnaeus*. **E.** *ezh2* expression in *N. furzeri* embryos (GRZ strain) during pre-diapause (onset of heartbeat), diapause escape (non-diapause development) and 3 days, 6 days and 1 month in diapause. **F, G.** *ezh2* expression during diapause and non-diapause development in *N. furzeri* (GRZ strain, F) and (MZM-0403 strain, G) from an independent study. **H.** *ezh2* expression during diapause and post-diapause development (4 days after exit from diapause) in *A. limnaeus*. **I, J.** *ezh1* and *ezh2* expression in *A. limnaeus* embryos in the trajectory leading to diapause entry (at 20°C) and non-diapause development (at 30°C). **K, L** ezh1 and ezh2 expression in *N. furzeri* tissues during aging. The expression levels of *ezh1* goes up with aging while the expression of *ezh2* decreases with age in brain, fat, heart, and muscle. Expression levels represent DE-Seq2 normalized counts. Values for panels A and E are on log scale. The P-values are from Wald’s test after multiple hypothesis correction using Benjamini Hochberg procedure (A-H). For longitudinal datasets, P-values were calculated using linear regression using SciPy linregress (I-L). All the datasets and P-values were downloaded from Killiverse.

Next, we looked at the bulk RNA-seq dataset during a developmental time-course including seven developmental stages ranging from dispersed cell stage to 24-somites at two different temperatures (20 °C and 30 °C) from the South American killifish *A. limnaeus*. The higher temperature (30 °C) leads to a continuous development trajectory, and the lower temperature (20 °C) will lead to all the embryos entering diapause [43]. In the diapause trajectory, *ezh1* is gradually upregulated showing a significant positive correlation leading to the critical window of diapause entry (24-somites stage), while *ezh2* is significant downregulated (Figures 2I, 2J and Figure S4). This expression pattern is reversed in the continuous development trajectory, with *ezh2* shows a significant positive correlation (Figures 2I, 2J and Figure S4). This suggests that the *ezh2*-to-*ezh1* switch in PRC2 complex is associated with diapause entry across diverse killifish species that likely evolved diapause independently. Interestingly, this switch occurs gradually during pre-diapause development even before the onset of diapause in response to temperature changes.

We next asked how the expression patterns of *ezh1* and *ezh2* changes with adult aging using a multi-tissue transcriptomic aging study available in Killiverse [44]. The *ezh1-*to*-ezh2* switch in PRC2 reverses during aging in the short-lived GRZ strain of *N. furzeri*: *ezh1* is significantly upregulated and *ezh2* significantly downregulated with age across multiple tissues including brain, fat, heart, and muscle (Figure 2K, 2L and Figure S5). This pattern is also recapitulated in another independent study on a slightly longer-lived MZM-04/10 strain during the brain aging (Figure S6) [45]. These observations suggest a shared epigenetic mechanism might underly both diapause and aging in killifish. PRC2-complex-mediated H3K27me3 is also important for mammalian aging [46, 47], and a switch from *ezh2*-to-*ezh1* in the PRC2 complex has also been observed during aging in mice liver [48]. Our observations from Killiverse suggest that a switch from *ezh2*-to-*ezh1* may be a conserved aspect of aging across species. This cross-species, cross-study, cross-lifespan pattern was identified within a single session without additional data processing, demonstrating the value of the Killiverse platform for hypothesis generation.

### A conserved microglial vitamin B_12_ transport signature in vertebrate brain aging

The vitamin B_12_ transporter transcobalamin provides another example of how Killiverse can uncover conserved, cell-type-associated aging signatures. Transcobalamin, encoded by *TCN2* in humans, binds vitamin B_12_ and mediates its uptake into cells. Vitamin B_12_ is an essential cofactor for methionine synthase and methylmalonyl-CoA mutase, linking it to one-carbon metabolism, DNA methylation, nucleotide synthesis, fatty acid metabolism, and mitochondrial energy production [49]. Defects in vitamin B_12_ transport and cellular uptake can cause neurological abnormalities, and vitamin B_12_ deficiency has been associated with cognitive and neuropsychiatric disorders [50]. Vitamin B_12_ has also been shown to directly regulate microglial transcriptional and metabolic states during neuroinflammation [51]. However, whether transcobalamin-mediated vitamin B_12_ transport changes during normal brain aging remains poorly understood.

Using Killiverse, we identified *LOC107394210*, the *N. furzeri* ortholog of human TCN2 (Figure S7), as an aging-associated gene in the killifish brain. Across independent bulk RNA-seq datasets, *LOC107394210* increased with age in both the GRZ (Figure 3A and Figures S8 and S9) and MZM-04/10 (Figure 3B and Figure S10) strains of *N. furzeri*, as well as in a second killifish species, *N. guentheri* Zanzibar Tan 14-02 (Figure 3C, 3D and Figure S11) [45, 52, 53]. Whole-lysate proteomics data also showed a concordant age-associated increase, but the difference is not statistically significant (Figure 3E and Figure S12) [54]. Single-nucleus RNA-seq localized *LOC107394210* expression predominantly to microglia (Figure 4A, 4B, 4C and Figures S13–S15).

**Figure 3.**
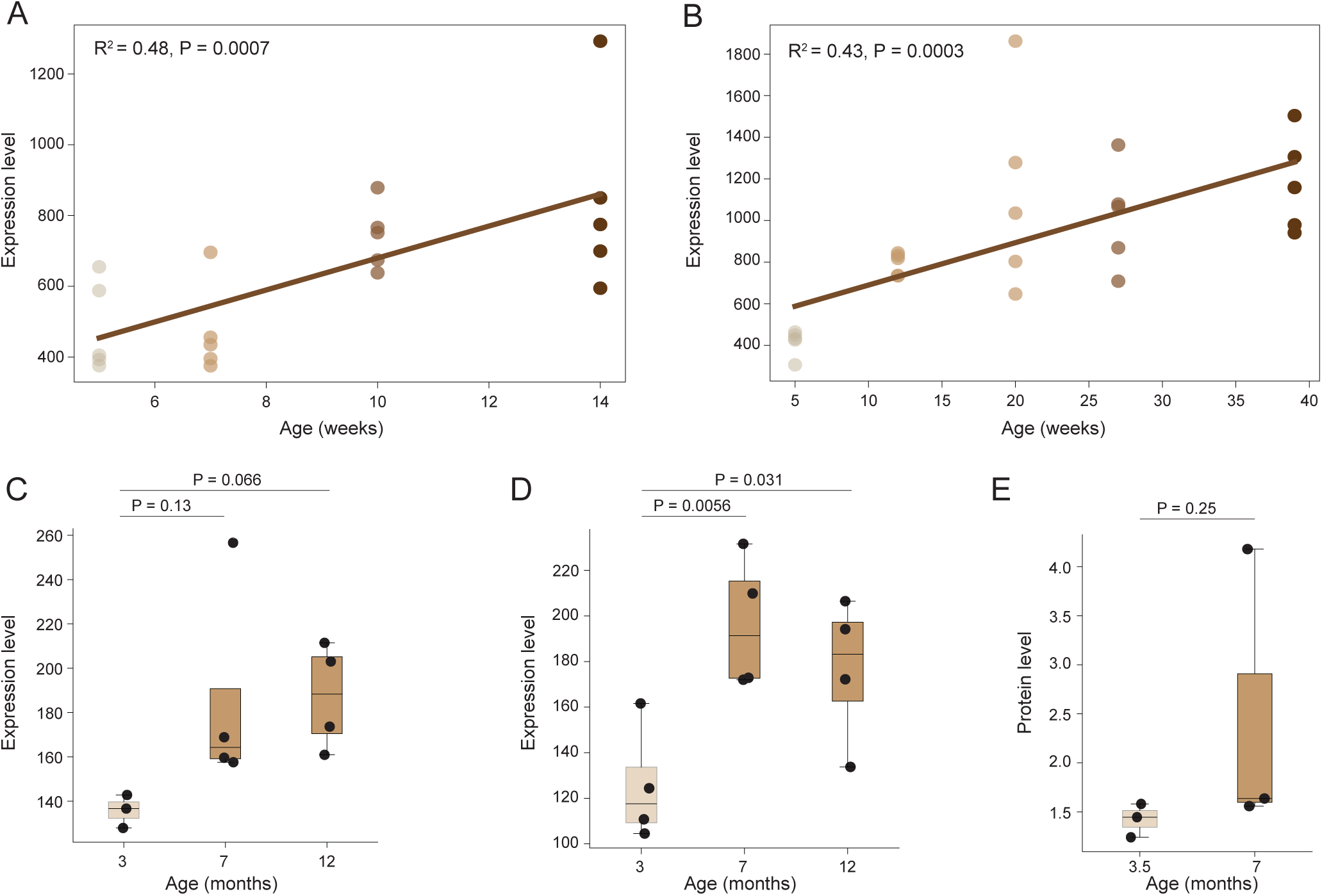
A conserved microglial vitamin B_12_ transport signature in vertebrate brain aging. **A, B**. Expression levels of the vitamin B_12_ transporter transcobalamin (*TCN2)* ortholog *LOC107394210* in the brain of male *N. furzeri* strains GRZ and MZM-04/10 goes significantly up with aging. **C, D.** Expression of *TCN2* orthologs also goes up in a longer-lived killifish species, *N. guentheri*, in both males (C) and females (D). **E.** Abundance of TCN2 protein from whole tissue lysate in the brains of male killifish *N. furzeri* (GRZ). Expression levels are DE-Seq2 normalized counts. For longitudinal datasets, P-values were calculated using linear regression using SciPy linregress. All the datasets and P-values were downloaded from Killiverse.

**Figure 4.**
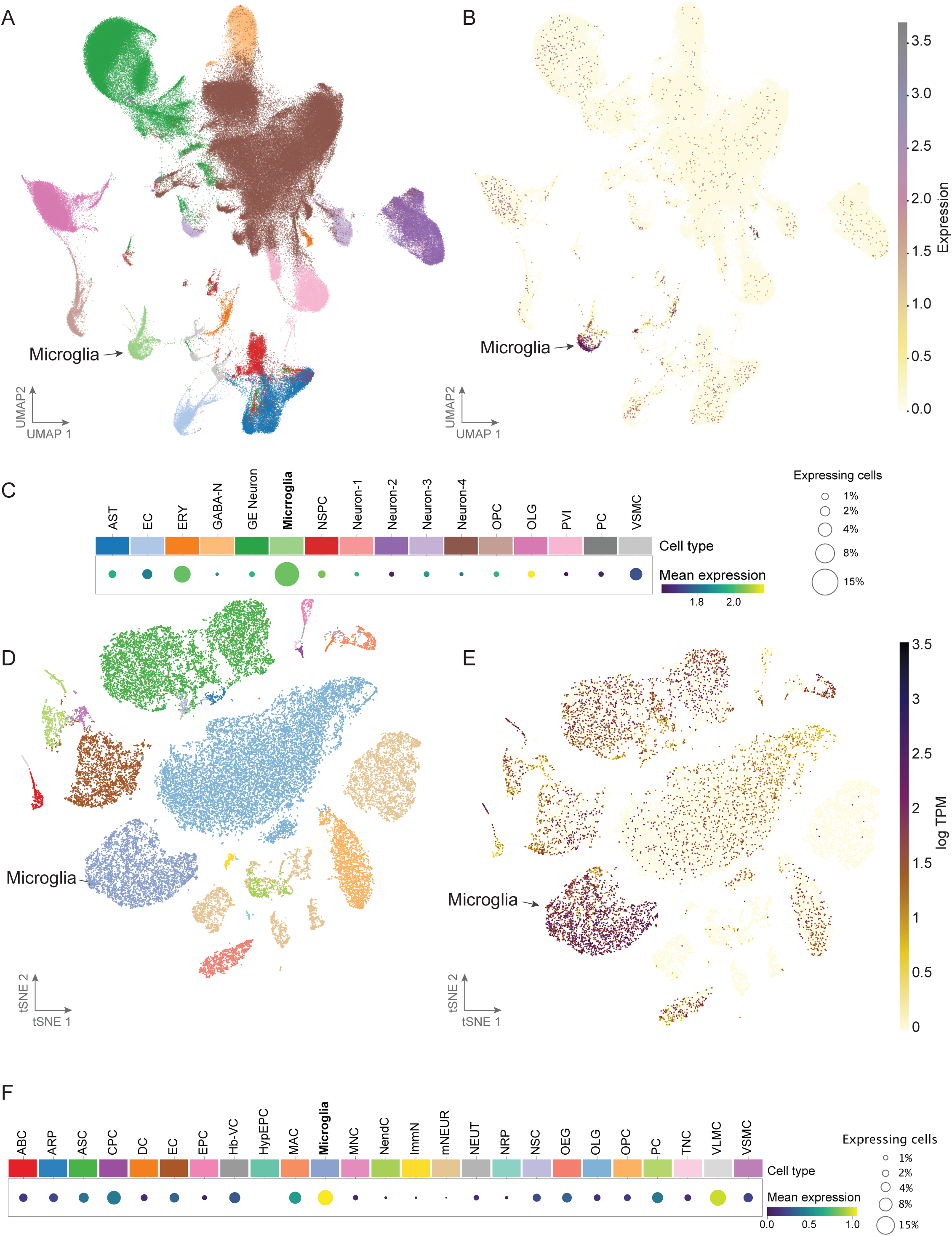
Expression of TCN2 in microglia. **A.** UMAP for the single-nucleus RNA-seq in the brain in N. furzeri colored by cell types. UMAP is from the original study available in KIlliverse. Males and female from two strains GRZ and ZMZ-1001 are plotted together. **B.** UMAP of single-nucleus RNA-seq data from the N. furzeri brain with cells colored by expression of the TCN2 ortholog, LOC107394210. **C.** Bubble plot with mean expression of LOC107394210 by cell types in N. furzeri with the highest expression in Microglia. See Table S3 for all cell type abbreviation details. **D.** tSNE plot for the single-cell RNA-seq in the brain in male mouse colored by cell types. The tSNE plots represents author’s original annotation and clustering, obtained from the Single Cell Portal at Broad Institute. **E.** tSNE plot for the single-cell RNA-seq in the brain in male mouse with cells colored by the expression of the Tcn2. **F.** Bubble plot with mean expression of LOC107394210 ortholog in mice across cell types in male mouse with the highest expression in Microglia. See Table S3 for all cell type abbreviation details.

To determine whether this cell-type specificity extends beyond killifish, we analyzed the aging mouse brain single-cell RNA-seq dataset [55]. We again observed expression of *Tcn2* predominantly in microglia (Figure 4D, 4E, and 4F). Age-associated changes in lipid metabolism, mitochondrial function, and inflammatory signaling are key features of aging microglia [56–58]. Because vitamin B_12_ is closely linked to mitochondrial and one-carbon metabolism, the conserved increase in *TCN2* expression observed here raises the possibility that regulation of cellular vitamin B_12_ availability may contribute to age-associated microglial dysfunction. Together, these analyses nominate transcobalamin-mediated vitamin B_12_ transport as a previously underappreciated component of vertebrate brain aging and highlight a conserved microglial expression program observed across multiple killifish strains, species, and independent bulk transcriptomics, proteomics, and single-cell datasets, illustrating how Killiverse can generate biologically meaningful hypotheses from heterogeneous multi-omics data.

### Fibrillin genes as conserved markers of vertebrate skin aging

The fibrillin family provides another example of how Killiverse can reveal conserved aging signatures across species. Fibrillins are large extracellular matrix glycoproteins that assemble into microfibrils, which provide mechanical support and contribute to tissue integrity and elasticity [59]. Although mutations in FBN genes have been linked to connective tissue disorders and aging-like phenotypes [60–62], the extent to which age-associated changes in fibrillin gene expression are conserved across vertebrate species remains unclear.

Using Killiverse, we observed a progressive decline in the expression of *fbn1* and *fbn3* in skin tissue with age in both GRZ and MZM-04/10 strains of *N. furzeri* (Figure 5A and 5B). A similar decline in *Fbn1* and *Fbn2* expression was also observed in aging mouse skin (Figure 5C). Given the emerging recognition of extracellular matrix remodeling as a hallmark of aging [63, 64], this conserved pattern suggests that reduced fibrillin expression may be a common feature of vertebrate skin aging and demonstrates the utility of Killiverse for identifying shared biological signals across independent datasets and species.

**Figure 5.**
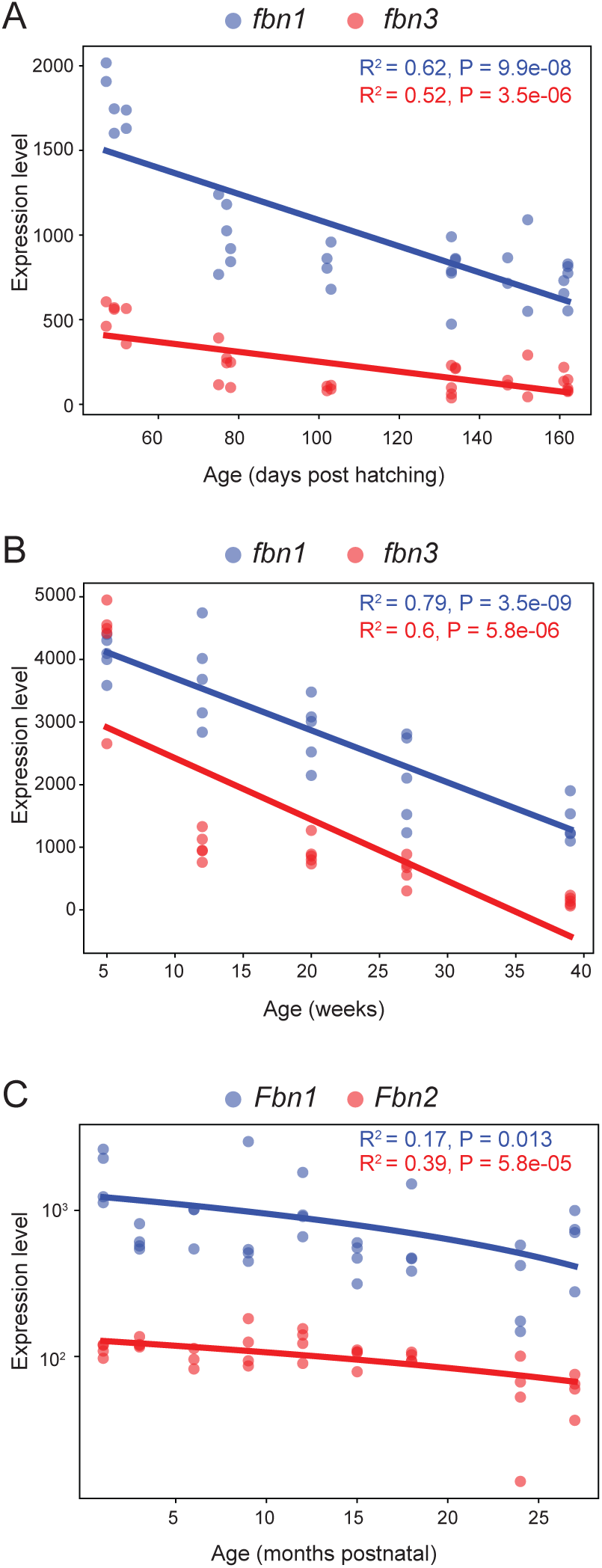
Fibrillin genes as conserved markers of vertebrate skin aging. **A, B**. Expression of the killifish fibrillin genes (*fbn1* and *fbn3*) in skin tissue during aging in the males of GRZ (**A**) and MZM-04/10 (**B**) strains. **C**. Expression of the fibrillin genes (*Fbn1* and *Fbn2*) during skin aging in male mice. Expression levels are DE-Seq2 normalized counts. For longitudinal datasets, P-values were calculated using linear regression using SciPy linregress.

## Conclusion

Killiverse provides a web-based platform for interactive exploration of multi-omics datasets, integrating bulk transcriptomic, single-cell, proteomic, and lipidomic measurements across studies, tissues, and species. By enabling rapid comparison of diverse datasets and experimental contexts, the platform facilitates discovery of conserved biological patterns and generation of new hypotheses. Through integration of ortholog mapping and cross-species datasets, Killiverse further enables findings from killifish studies to be placed within a broader vertebrate context.

Current limitations include reliance on processed data objects for certain modalities and incomplete harmonization of annotations across single cell studies. As the community grows and new datasets emerge, we aim to expand Killiverse into a comprehensive community resource for killifish genomics, with continued incorporation of additional modalities and improved cross-species and cross-study mapping.

## Availability and requirements

Project name: Killiverse

Project home page: https://killiverse.org/

Operating system(s): Platform independent

Other requirements: modern browser (Apple Safari, Google Chrome, Mozilla Firefox, or Microsoft Edge)

License: Creative Commons Attribution 4.0 International

## Availability of data

The code underlying Killiverse are available on GitHub (https://github.com/SinghLabUCSF/Killiverse). Details of the included studies and datasets are provided in Supplementary Table S1.

## Author contributions

PPS conceived the study, performed part of the analyses and contributed to the manuscript writing. AM performed the software development and analysis, prepared figures and tables, and drafted the manuscript. All authors read and approved the final manuscript.

## Acknowledgements

We thank Anastasia Paulmann, Dario Valenzano, and Itamar Harel for sharing their Seurat data objects, Bérénice Benayoun for sharing the code to process human GTEx data, Rogelio Barajas for contributing graphics for the website, and all the members of Singh lab for contributing ideas in Killiverse development and comments on the manuscript.

