## Supplementary_Data for "Killiverse: an interactive multi-omics web resource for killifish"

**Table S1: Studies with links to Killiverse, GEO accession numbers for the datasets, assay type, scientific name of the species and strain**

| No. | Study title with hyperlink to Killiverse | GEO accession of dataset | RNA-seq modality | Species (strain) |
|---|---|---|---|---|
| 1. | Multi-tissue transcriptomic aging atlas reveals predictive aging biomarkers in the killifish | PRJNA1274512 | bulk RNA-seq (6 age groups) | *N. furzeri* (GRZ) |
| 2. | Vertebrate diapause preserves organisms long term through Polycomb complex members | PRJNA503701 | bulk RNA-seq | *N. furzeri* (GRZ) |
| 3. | Widespread sex dimorphism across single-cell transcriptomes of adult African turquoise killifish tissues[a] | PRJNA952805 | single cell RNA-seq | *N. furzeri* (GRZ) |
| 4. | Transcriptomes of aging brain, heart, muscle, and spleen from female and male African turquoise killifish | SRP430823 (PRJNA952180) | bulk RNA-seq | *N. furzeri* (GRZ) |
| 5. | The killifish germline regulates longevity and somatic repair in a sex-specific manner | PRJNA1045681 | bulk RNA-seq | *N. furzeri* (GRZ) |
| 6. | Regulation of life span by the gut microbiota in the short-lived African turquoise killifish | PRJNA379208 | bulk RNA-seq | *N. furzeri* (GRZ) |
| 7. | An automated feeding system for the African killifish reveals effects of dietary restriction on lifespan and allows scalable assessment of associative learning | GSE216369 (PRJNA893228) | bulk RNA-seq | *N. furzeri* (GRZ) |
| 8. | Refeeding-associated AMPKγ1 complex activity is a hallmark of health and longevity | PRJNA817434 | bulk RNA-seq | *N. furzeri* (GRZ) |
| 9. | Genetic perturbation of AMP biosynthesis extends lifespan and restores metabolic health in a naturally short-lived vertebrate | GSE190757 (PRJNA788399) | bulk RNA-seq | *N. furzeri* (GRZ) |
| 10. | RNA-seq of the aging brain in the short-lived fish N. furzeri - conserved pathways and novel genes associated with neurogenesis | GSE52462 (PRJNA229052) | bulk RNA-seq (5 age groups) | *N. furzeri* (MZM-04/10) |

| | | | | |
|---|---|---|---|---|
| 11. | Evolution of diapause in the African turquoise killifish by remodeling the ancient gene regulatory landscape (Dataset 1) | GSE185817 (PRJNA770943) | bulk RNA-seq | *N. furzeri* (GRZ) |
| 12. | Evolution of diapause in the African turquoise killifish by remodeling the ancient gene regulatory landscape (Dataset 2) | Supplemental information S7 | lipidomics | *N. furzeri* (GRZ) |
| 13. | Evolution of diapause in the African turquoise killifish by remodeling the ancient gene regulatory landscape (Dataset 3) | Supplemental information S7 | lipidomics | *N. furzeri* (GRZ) *A. striatum* (Aquarium) |
| 14. | MicroRNA miR-29 controls a compensatory response to limit neuronal iron accumulation during adult life and aging | GSE79825 (PRJNA317074) | bulk RNA-seq | *N. furzeri* (MZM-04/10) |
| 15. | Reduced proteasome activity in the aging brain results in ribosome stoichiometry loss and aggregation | GSE150318 (PRJNA631760) | bulk RNA-seq | *N. furzeri* (MZM-04/10) |
| 16. | Aging triggers H3K27 trimethylation hoarding in the chromatin of Nothobranchius furzeri skeletal muscle | GSE135032 (PRJNA557199) | bulk RNA-seq | *N. furzeri* (MZM-04/10) |
| 17. | Dynamic regulation of gonadal transposon control across the lifespan of the naturally short-lived African turquoise killifish | PRJNA854614 | bulk RNA-seq | *N. furzeri* (GRZ) |
| 18. | Circular RNAs in the ageing African turquoise killifish | GSE134065 (PRJNA553674) | bulk RNA-seq | *N. furzeri* (GRZ) |
| 19. | The effect of meclofenoxate on the transcriptome of aging brain of Nothobranchius guentheri annual killifish | PRJNA779252 | bulk RNA-seq | *N. guentheri* (Zanzibar Tan 14-02) |
| 20. | Exercise datasets[b] | PRJNA407099 (muscle), PRJNA407101 (skin), PRJNA407084 (brain), PRJNA407091 (liver) | bulk RNA-seq | *N. furzeri* (MZM-04/10) |

| | | | | |
|---|---|---|---|---|
| 21. | Longitudinal RNA-seq analysis of vertebrate aging identifies mitochondrial complex I as a small-molecule-sensitive modifier of lifespan (Dataset I) | GSE66712 (PRJNA277768) | bulk RNA-seq | *N. furzeri* (MZM-04/10) |
| 22. | Longitudinal RNA-seq analysis of vertebrate aging Identifies mitochondrial complex I as a small-molecule-sensitive modifier of lifespan (Dataset II) | GSE66712 (PRJNA277768) | bulk RNA-seq (5 age groups) | *N. furzeri* (MZM-04/10) |
| 23. | RNAseq analysis of brain aging in wild specimens of short-lived turquoise killifish: Commonalities and differences with aging under laboratory conditions (Dataset I) | GSE183039 (PRJNA758914) | bulk RNA-seq | *N. furzeri* (MZM-04/10) |
| 24. | RNAseq Analysis of Brain Aging in Wild Specimens of short-lived turquoise killifish: Commonalities and differences with aging under laboratory conditions (Dataset II) | GSE183039 (PRJNA758914) | bulk RNA-seq | *N. furzeri* (MZM-04/10), *N. furzeri* (MZM-A41), |
| 25. | Age-related dysregulation of the retinal transcriptome in African turquoise killifish | GSE255363 (PRJNA1074746) | bulk RNA-seq | *N. furzeri* (GRZ) |
| 26. | Changes in regeneration-responsive enhancers shape regenerative capacities in vertebrates | PRJNA559885 | bulk RNA-seq | *N. furzeri* (GRZ) |
| 27. | Analysis of different strains of the turquoise killifish *Nothobranchius furzeri* identifies transcriptomic signatures associated with heritable lifespan differences | GSE103132 (PRJNA400236) | bulk RNA-seq (4 age groups) | *N. furzeri* (GRZ) |
| 28. | Temperature-dependent vitamin D signaling regulates developmental trajectory associated with diapause in an annual killifish [c] | PRJNA272154 | bulk RNA-seq | *A. limnaeus* (Quisiro) |
| 29. | Single housing of juveniles accelerates early-stage growth but extends adult lifespan in African turquoise killifish (Dataset I) | GSE245483 (PRJNA1028576) | bulk RNA-seq (5 age groups) | *N. furzeri* (GRZ) |
| 30. | Single housing of juveniles accelerates early-stage growth but extends adult lifespan in African turquoise killifish (Dataset II) | GSE245483 (PRJNA1028576) | bulk RNA-seq (5 age groups) | *N. furzeri* (GRZ) |

| | | | | |
|---|---|---|---|---|
| 31. | Single housing of juveniles accelerates early-stage growth but extends adult lifespan in African turquoise killifish (Dataset III) | GSE245483 (PRJNA1028576) | bulk RNA-seq | *N. furzeri* (GRZ) |
| 32. | The killifish germline regulates longevity and somatic repair in a sex-specific manner (Dataset I) | GSE248741 (PRJNA1045681) | single cell RNA-seq | *N. furzeri* (GRZ) |
| 33. | The killifish germline regulates longevity and somatic repair in a sex-specific manner (Dataset II) | GSE248741 (PRJNA1045681) | single cell RNA-seq | *N. furzeri* (GRZ) |
| 34. | The killifish germline regulates longevity and somatic repair in a sex-specific manner (Dataset III) | GSE248741 (PRJNA1045681) | single cell RNA-seq | *N. furzeri* (GRZ) |
| 35. | The killifish germline regulates longevity and somatic repair in a sex-specific manner (Dataset IV) | GSE248741 (PRJNA1045681) | single cell RNA-seq | *N. furzeri* (GRZ) |
| 36. | The genome of *Austrofundulus limnaeus* offers insights into extreme vertebrate stress tolerance and embryonic development | PRJNA272154 | bulk RNA-seq | *A. limnaeus* (Quisiro) |
| 37. | Spontaneous onset of cellular markers of inflammation and genome instability during aging in the immune niche of the naturally short-lived turquoise killifish (*Nothobranchius furzeri*) | BioStudies S-BSST2265 | single cell RNA-seq | *N. furzeri* (GRZ) |
| 38. | Sodium-glucose co-transporter 2 inhibition improves age-dependent kidney microvascular rarefaction | PRJNA1265492 | single nucleus RNA-seq | *N. furzeri* (GRZ) |
| 39. | Tissue-specific landscape of protein aggregation and quality control in an aging vertebrate (Dataset 1) | Supplemental information Table S2 | proteomics | *N. furzeri* (GRZ) |
| 40. | Tissue-specific landscape of protein aggregation and quality control in an aging vertebrate (Dataset 2) | Supplemental information Table S2 | proteomics | *N. furzeri* (GRZ) |
| 41. | Tissue-specific landscape of protein aggregation and quality control in an aging vertebrate (Dataset 3) | Supplemental information Table S2 | proteomics | *N. furzeri* (GRZ) |

| | | | | |
|---|---|---|---|---|
| 42. | A multi-omic atlas in the African turquoise killifish reveals increased glucocorticoid signaling as a hallmark of brain aging [d] | PRJNA1105049 | single nucleus RNA-seq | *N. furzeri* (GRZ), *N. furzeri* (ZMZ-1001) |
| 43. | Insights into Sex Chromosome Evolution and Aging from the Genome of a Short-Lived Fish | PRJEB5837 | bulk RNA-seq | *N. furzeri* (GRZ), *N. furzeri* (MZM-0403) |
| 44. | The human and non-human primate developmental GTEx projects [e] | dbGaP accession phs000424 (v10.p2) | bulk RNA-seq (6 age groups) | *Homo sapiens* |
| 45. | Ageing hallmarks exhibit organ-specific temporal signatures | GSE132040 (PRJNA544748) | bulk RNA-seq (10 age groups) | *Mus musculus* |

a. Seurat object available here: https://figshare.cosm/articles/dataset/Annotated_Seurat_object_containing_cells_for_a_single-cell_transcriptomic_atlas_of_adult_female_and_male_African_turquoise_killifish_/22766894
b. Study not published
c. Processed using GCF_001266775.1 *A. limnaeus* genome assembly.
d. Seurat object available here: https://figshare.com/articles/dataset/Annotated_Seurat_object_for_a_single-nuclei_transcriptomic_atlas_of_aging_female_and_male_African_turquoise_killifish_brains_of_the_GRZ_and_ZMZ-1001_strains/31847155?file=63065740
e. Raw count files and TPM files were obtained from gtexportal.org (https://gtexportal.org/home/downloads/adult-gtex/bulk_tissue_expression).

**Ortholog mapping using reciprocal best BLAST hits**

**Table S2. List of species and the genome assemblies used for ortholog identification.**

| Common name | Scientific name | Genome assembly |
|---|---|---|
| Human | *Homo sapiens* | GCF_000001405.40 |
| Mouse | *Mus musculus* | GCF_000001635.27 |
| Zebrafish | *Danio rerio* | GCF_049306965.1 |
| Japanese rice fish | *Oryzias latipes* | GCF_002234675.1 |
| African turquoise killifish | *Nothobranchius furzeri* | GCF_001465895.1 |
| Chicken | *Gallus gallus* | GCF_016699485.2 |
| Rat | *Rattus norvegicus* | GCF_036323735.1 |
| South American killifish | *Austrofundulus limnaeus* | GCF_001266775.1 |

Protein FASTA files were obtained from NCBI for the reference genome assemblies listed above. FASTA files were filtered to retain the longest protein sequence for each gene. BLASTP (version 2.16.0+) was run twice for every species. First, *N. furzeri's* protein FASTA file was used as the subject and the other species' protein FASTA file as the query. In the second run, *N. furzeri's* protein FASTA file was used as the query and the other species' protein FASTA file as the subject. E-value threshold was set to 0.01. BLASTP output files containing results for protein-protein sequence similarity searches were filtered to retain only the top hits (lowest e-value) for each gene. If both *N. furzeri* and the other species shared a pair of genes as the top hit, then they were considered orthologs. If no bidirectional hit was found, the top BLASTP hit obtained with *N. furzeri* as the query against other species was treated as the ortholog. Orthologs were identified for 22,119 out of 22,207 protein coding genes of *N. furzeri* in at least one of the other species.

**Table S3. Full names of all the cell types in killifish and brain from Figure 4.**

| Mouse cells | | Killifish cells | |
|---|---|---|---|
| **Abbreviation** | **Full name** | **Abbreviation** | **Full name** |
| ABC | Arachnoid barrier cells | AST | Astrocytes radial glia |
| ARP | Astrocyte-restricted precursors | EC | Ependymal cells |
| ASC | Astrocytes | ERY | Erythrocytes |
| CPC | Choroid plexus epithelial cells | GABA-N | GABAergic neurons |
| DC | Dendritic cells | GE Neuron | Granule excitatory neurons |
| EC | Endothelial cells | Microglia | Microglia |
| EPC | Ependymocytes | NSPC | NSPCs |
| Hb-VC | Hemoglobin-expressing vascular cells | Neuron-1 | Neurons misc 1 |
| HypEPC | Hypendymal cells | Neuron-2 | Neurons misc 2 |
| MAC | Macrophages | Neuron-3 | Neurons misc 3 |
| Microglia | Microglia | Neuron-4 | Neurons misc 4 |
| MNC | Monocytes | OPC | Oligodendrocyte progenitor cells |
| NendC | Neuroendocrine cells | OLG | Oligodendrocytes |
| ImmN | Immature neurons | PVI | PV interneurons |
| mNEUR | Mature neurons | PC | Purkinje cells |
| NEUT | Neutrophils | VSMC | Vascular smooth muscle cells |
| NRP | Neuronal restricted precursors | | |
| NSC | Neural stem cells | | |
| OEG | olfactory ensheathing glia | | |
| OLG | oligodendrocytes | | |
| OPC | oligodendrocyte precursor cells | | |
| PC | Pericytes | | |
| TNC | Tanycytes | | |
| VLMC | Vascular and leptomeningeal cells | | |
| VSMC | Vascular smooth muscle cells | | |

**Figure S1.** Killifish datasets filtered by library preparation protocol “Bulk RNA-seq” and condition “diapause”.

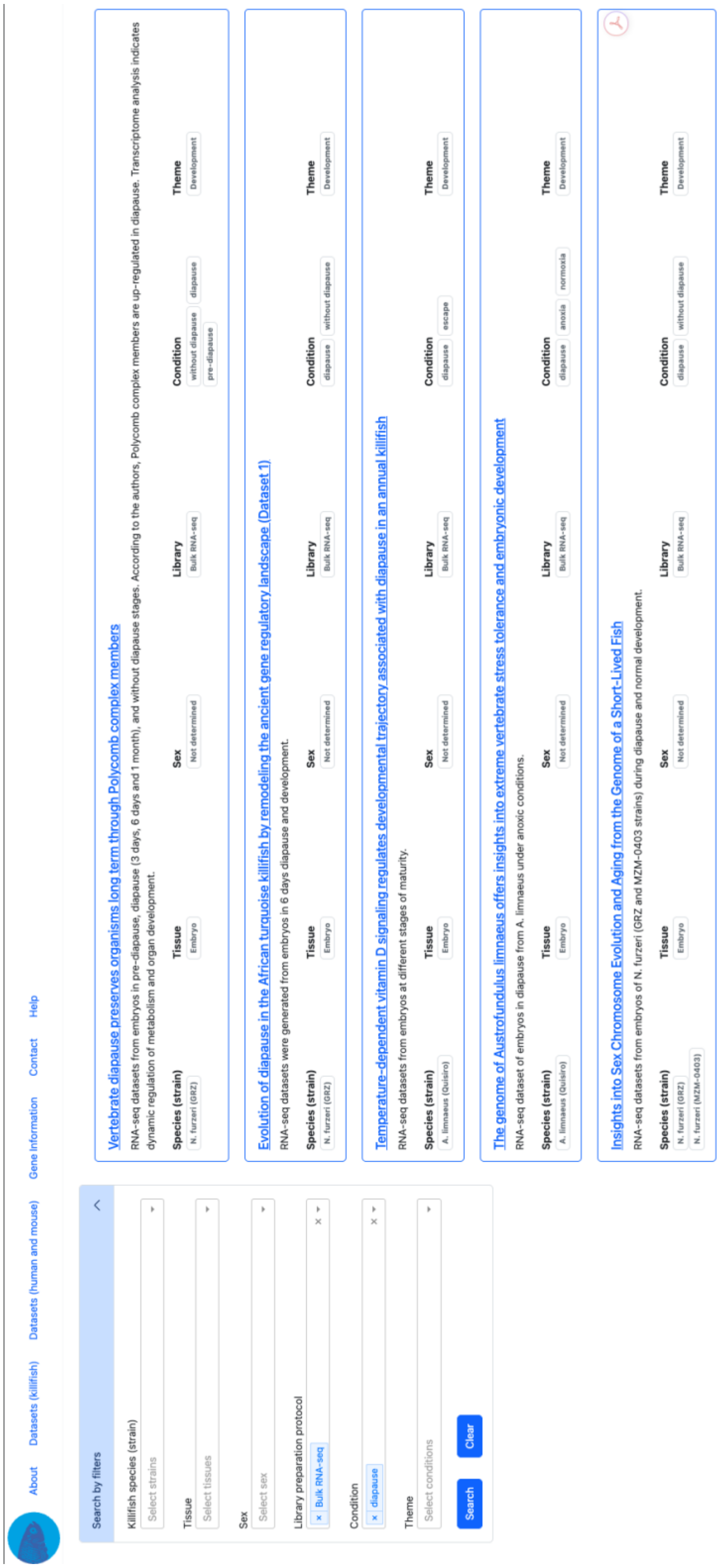

**Figure S2.** Box plot for the expression of *ezh1* and *ezh2* during pre-diapause (PreD), diapause 3-day (D3d), diapause 6-day (D6d), diapause 1-month (D1m) and non-diapause (NonD). Expression of *ezh1* is consistently higher in samples from diapause stage as compared to those in pre-diapause and non-diapause stages. The opposite is true for *ezh2*. This is part of the study [Vertebrate diapause preserves organisms long term through Polycomb complex members](#).

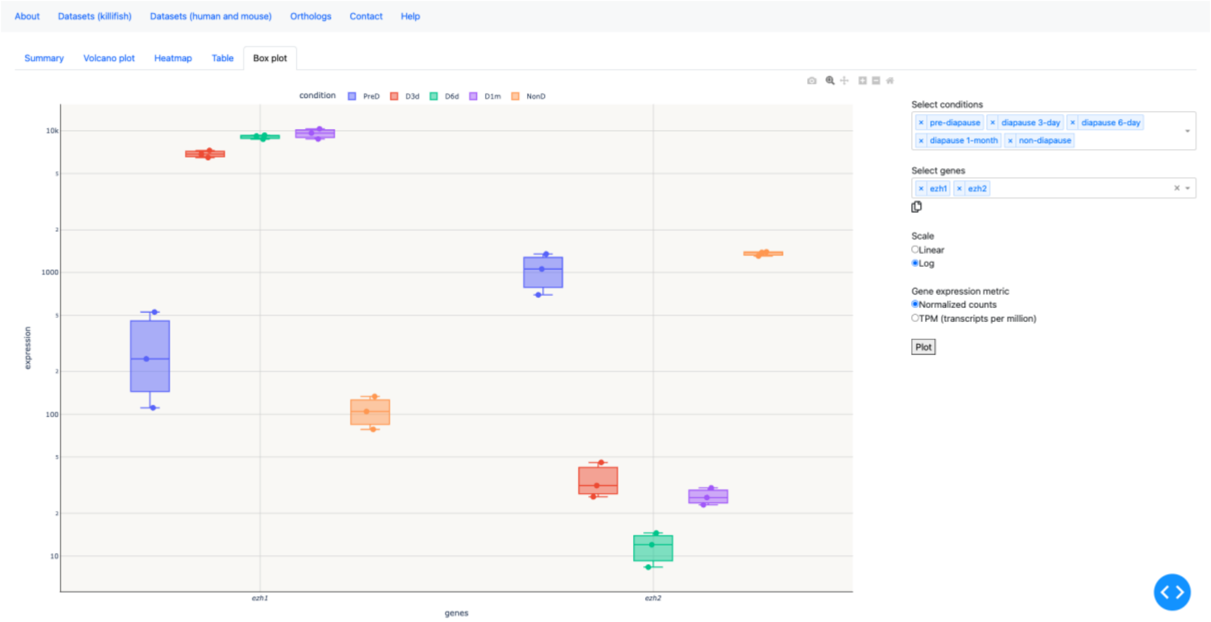

**Figure S3.** Heatmap for the expression of *ezh1* and *ezh2* genes during diapause (6 days) and development. Expression of *ezh1* tends to be higher in samples from diapause stage as compared to those development stage. The opposite is true for *ezh2*. This is part of the study [Evolution of diapause in the African turquoise killifish by remodeling the ancient gene regulatory landscape](#).

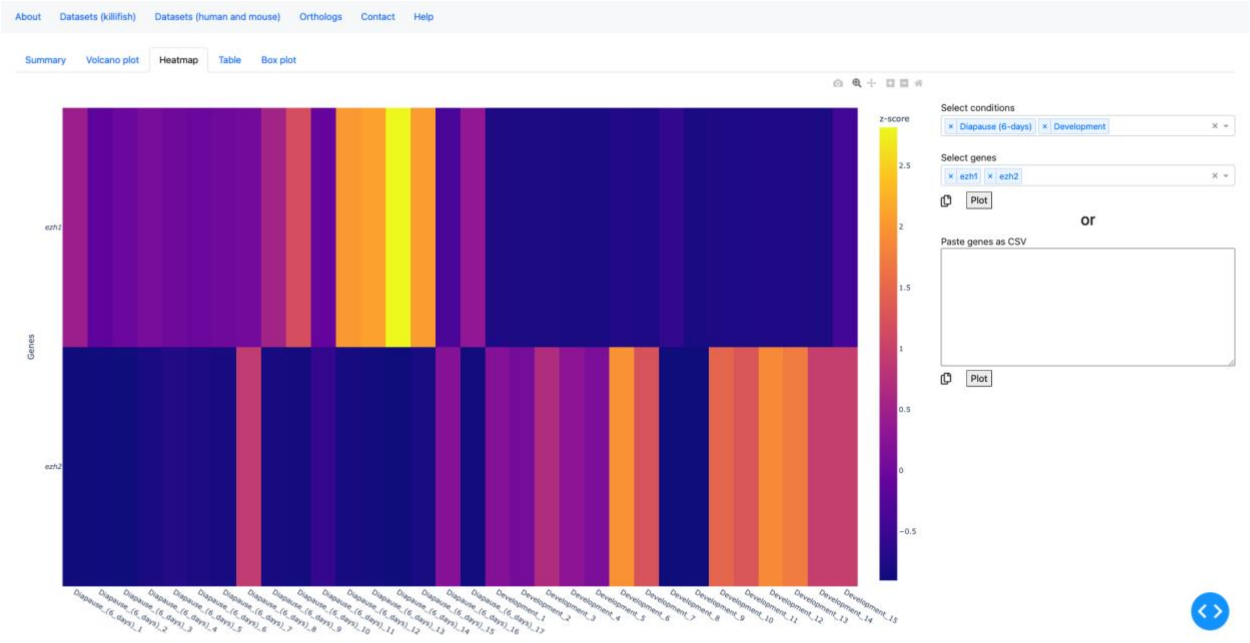

**Figure S4.** In the study titled Temperature-dependent vitamin D signaling regulates developmental trajectory associated with diapause in an annual killifish, embryos incubated at 20 °C committed to diapause trajectory while those incubated at 30 °C underwent normal development (escape trajectory). Expression of *ezh1* increased while that of *ezh2* declined as the embryo developed from dispersed cell stage (stage 1) to 24-pair somite stage (stage 7) during diapause. The opposite was true during normal development.

**A.** Expression of *ezh1* during diapause and normal development (escape).

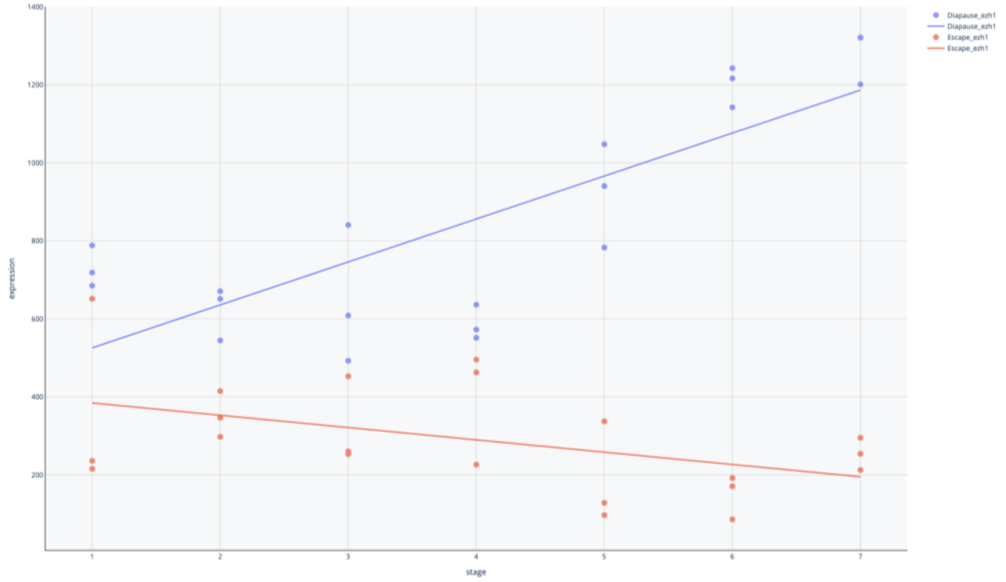


**B.** Expression of *ezh2* during diapause and normal development (escape).

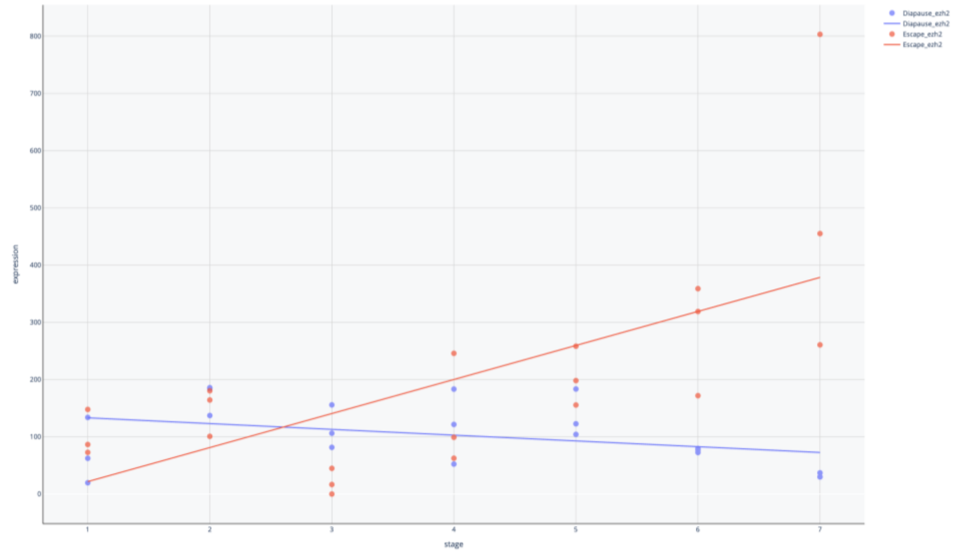

**Figure S5.** In the study titled Multi-tissue transcriptomic aging atlas reveals predictive aging biomarkers in the killifish, *ezh1* upregulated with age, while *ezh2* is downregulated across brain, fat, heart, and muscle.

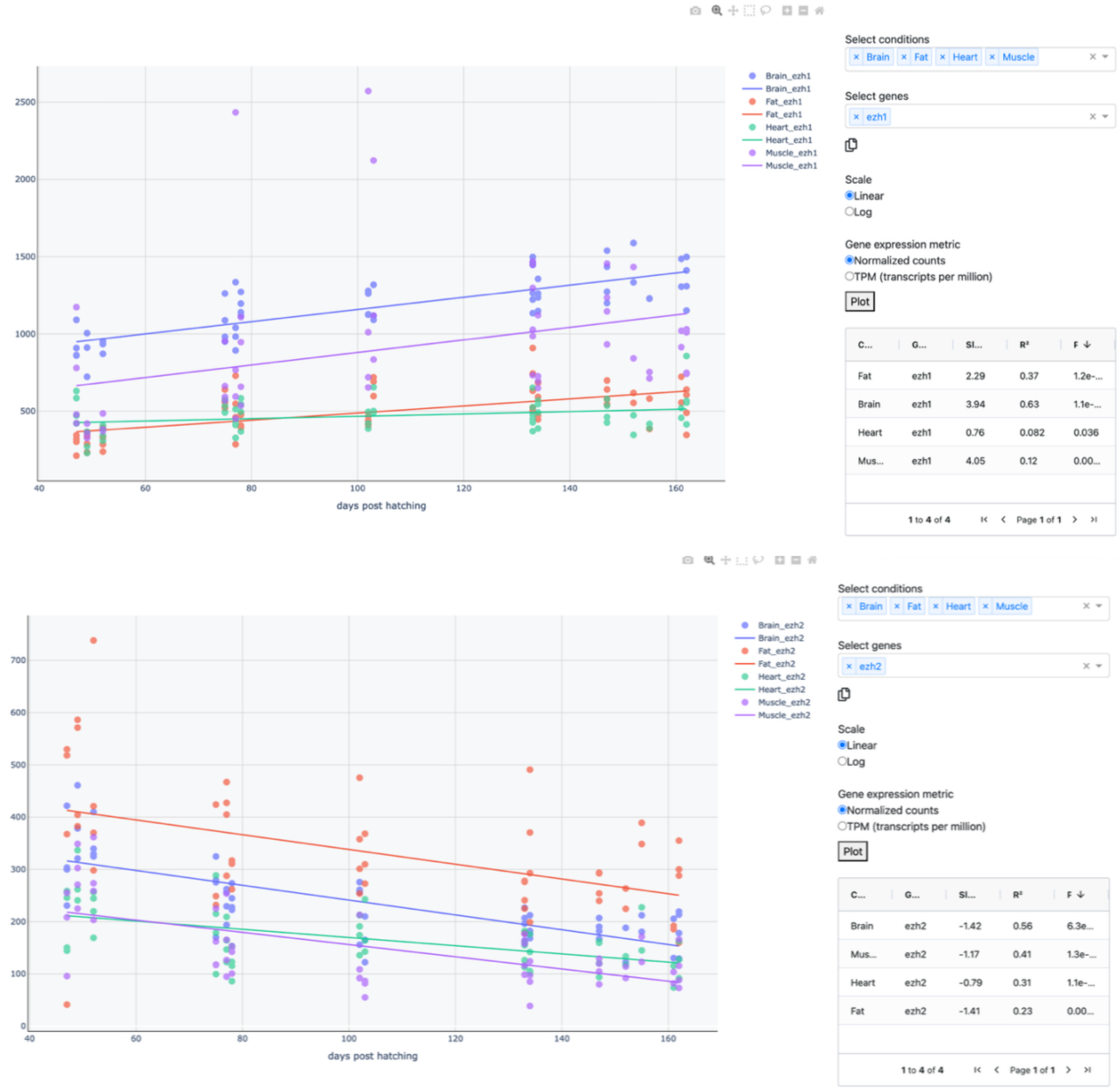

**Figure S6.** Line graph for the expression of *ezh1* and *ezh2* with age in male brain from *N. furzeri* (MZM-04/10) from study titled RNA-seq of the aging brain in the short-lived fish N. furzeri - conserved pathways and novel genes associated with neurogenesis

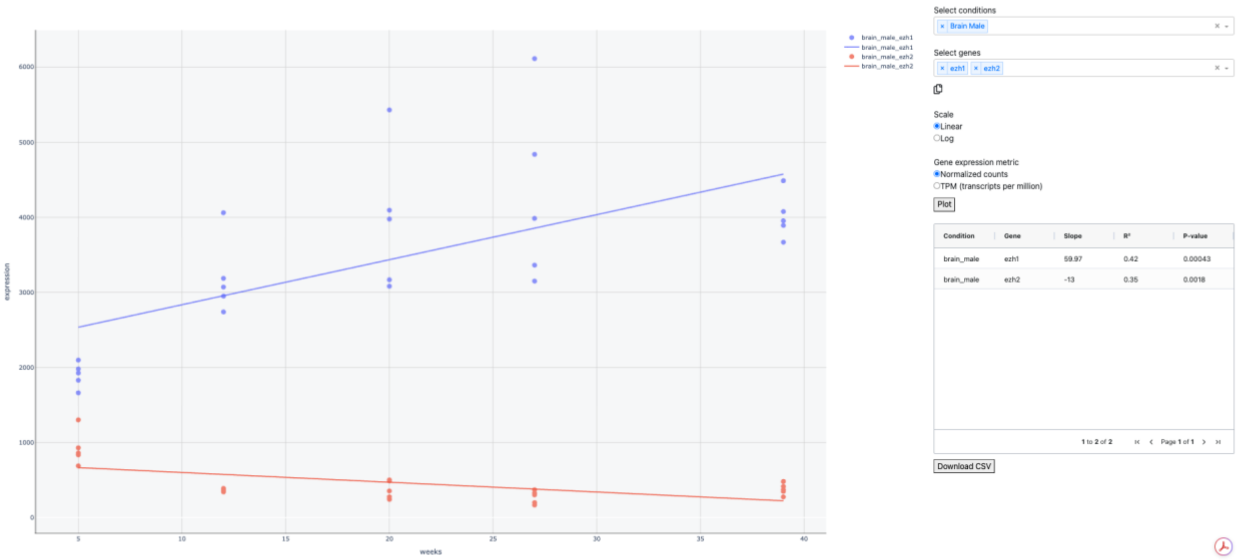

**Figure S7.** Information of *LOC107394210* gene in *N. furzeri* from Killiverse. This includes a brief description, GO annotations and links to NCBI Gene pages for orthologs from other model organisms.

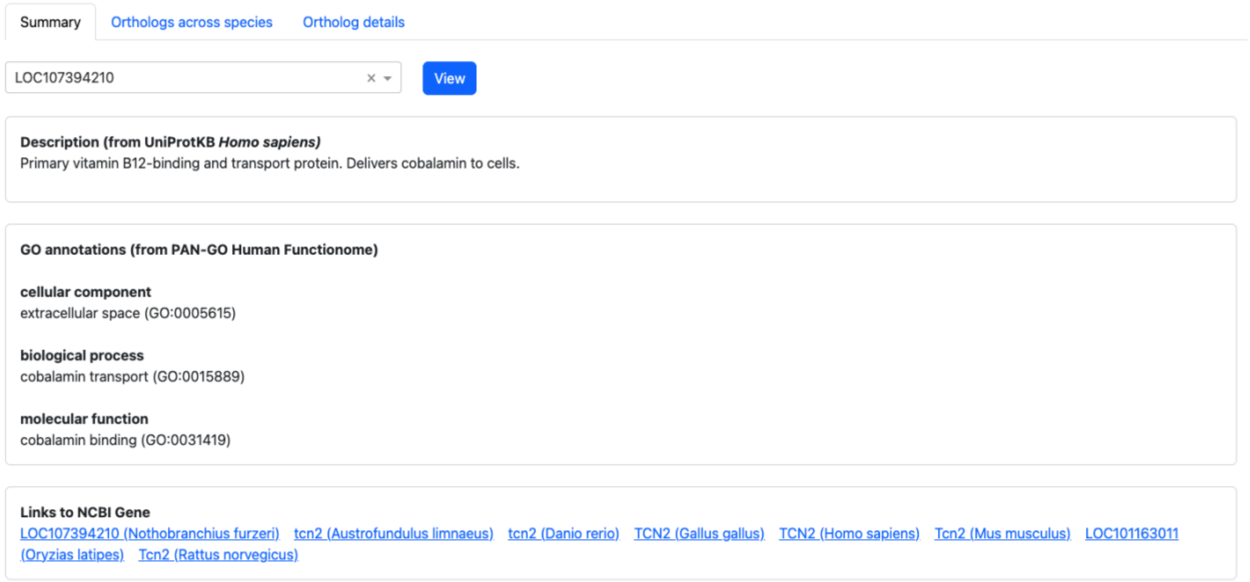

**Figure S8.** Line graph for the expression of *LOC107394210* with age in male brain from *N. furzeri* (GRZ) from study titled Analysis of different strains of the turquoise killifish Nothobranchius furzeri identifies transcriptomic signatures associated with heritable lifespan differences.

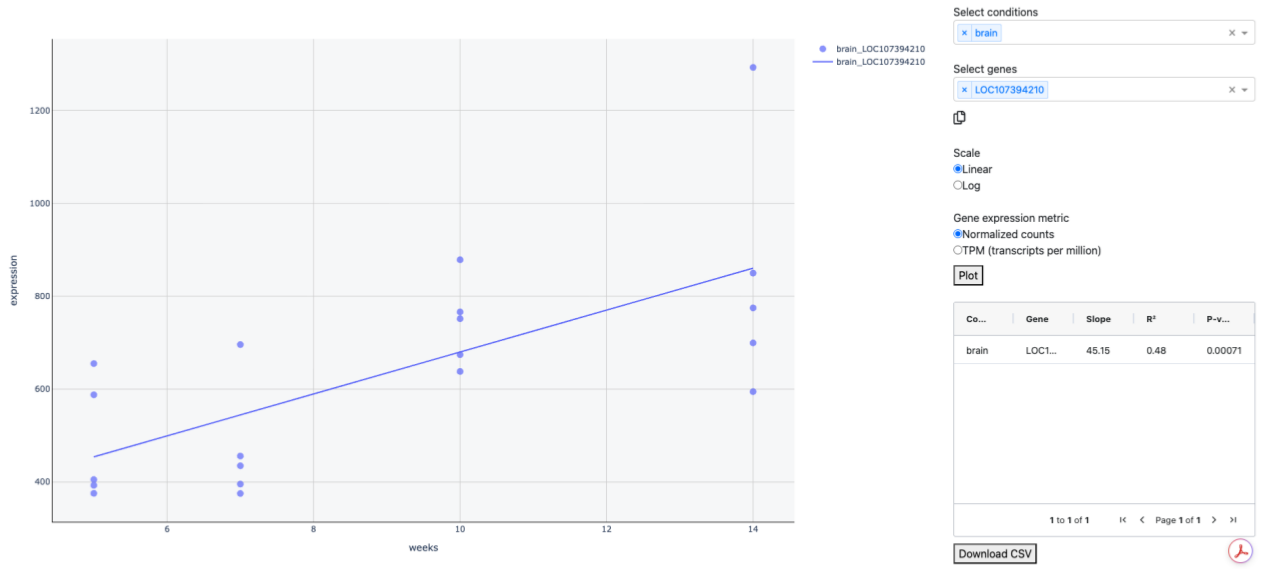

**Figure S9.** Spearman’s rank correlation and adjusted p-value with age for *LOC107394210* expression. This data was obtained from *N. furzeri* (GRZ) in study titled Multi-tissue transcriptomic aging atlas reveals predictive aging biomarkers in the killifish.

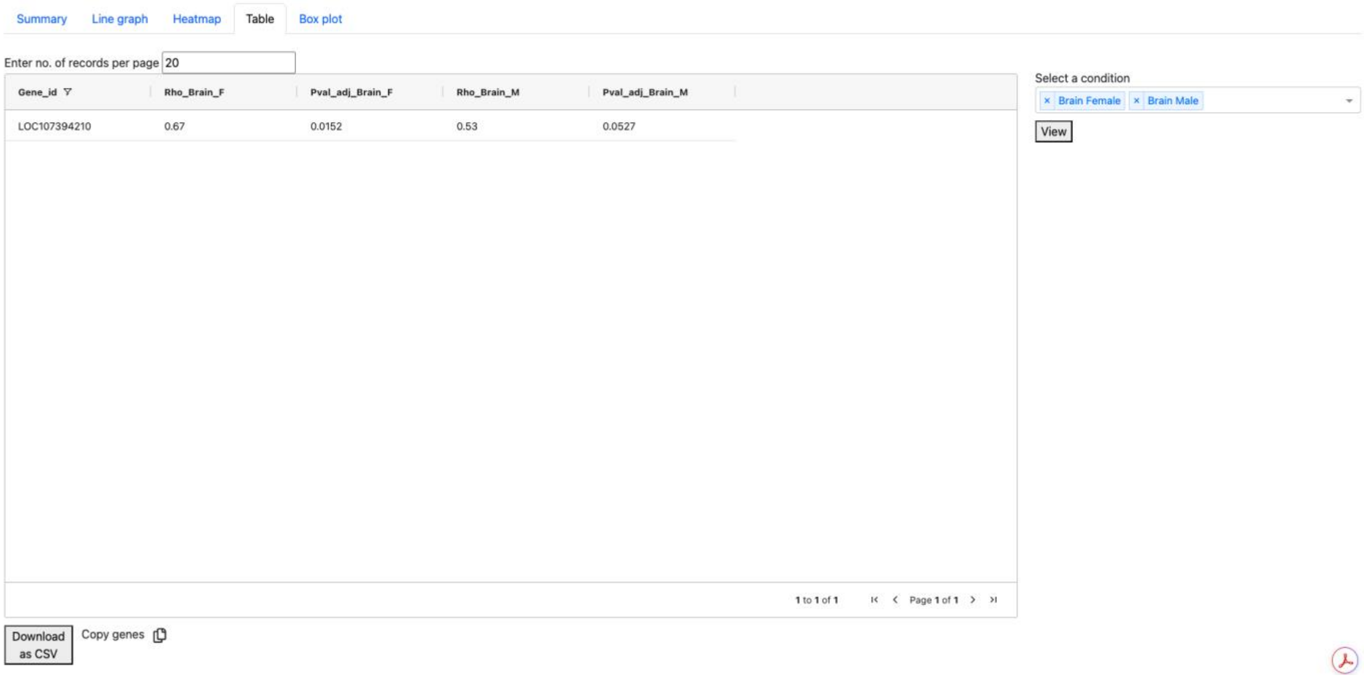


| Gene_id | Rho_Brain_F | Pval_adj_Brain_F | Rho_Brain_M | Pval_adj_Brain_M |
|---|---|---|---|---|
| LOC107394210 | 0.67 | 0.0152 | 0.53 | 0.0527 |

**Figure S10.** Line graph for the expression of *LOC107394210* with age in male brain from *N. furzeri* (MZM-04/10) from study titled RNA-seq of the aging brain in the short-lived fish N. furzeri - conserved pathways and novel genes associated with neurogenesis

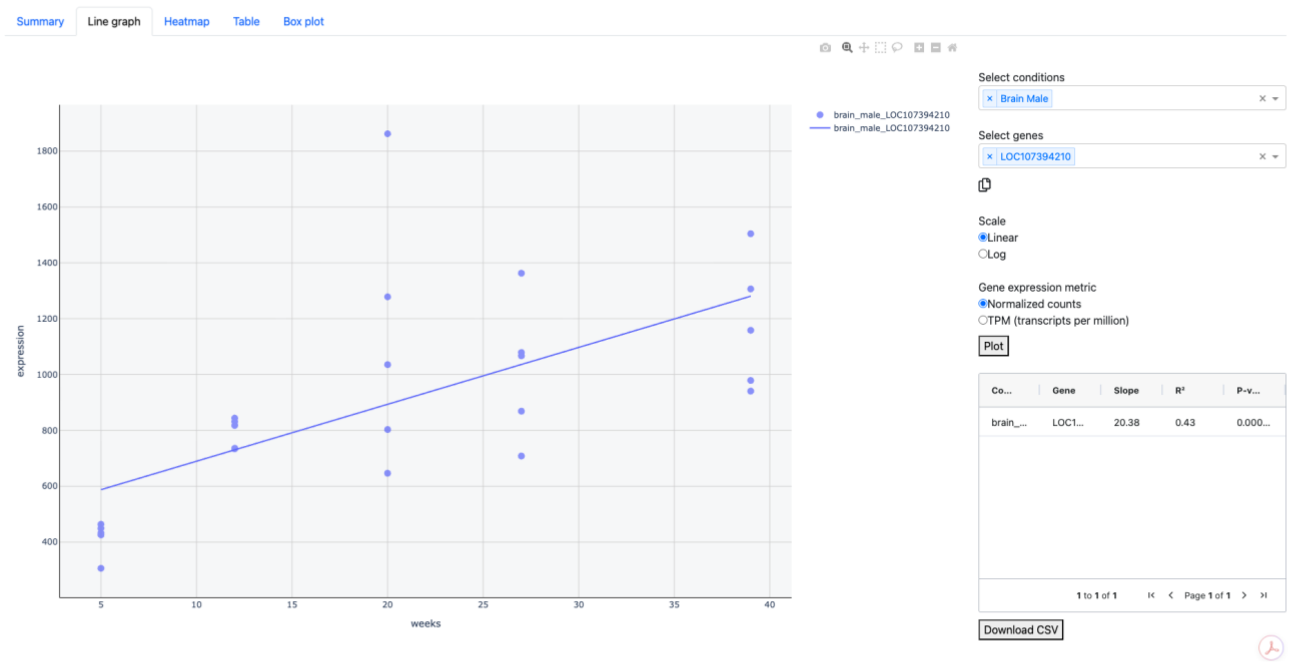


**Figure S11.** Box plot for the expression of *LOC107394210* with age in male and female brain from *N. guentheri* (Zanzibar TAN 14-02) from study titled The effect of meclofenoxate on the transcriptome of aging brain of Nothobranchius guentheri annual killifish

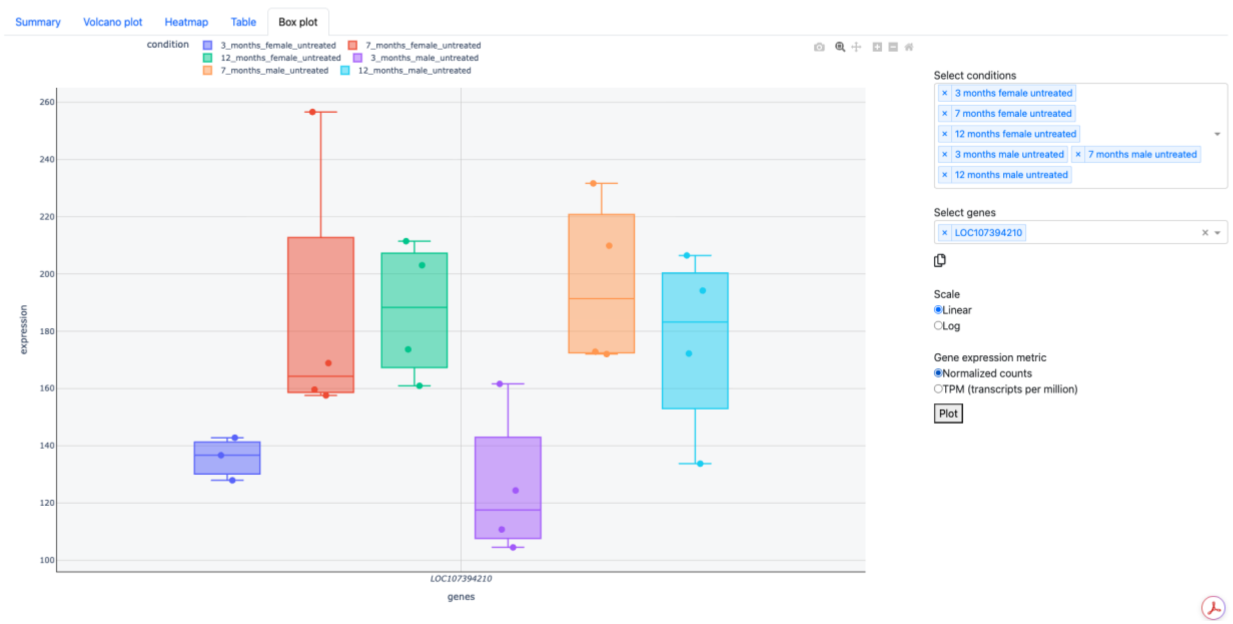

**Figure S12.** Expression of LOC107394210 in total lysate proteome data from young (3.5 months) and old (7 months) male brain of *N. furzeri* (GRZ), from the study Tissue-specific landscape of protein aggregation and quality control in an aging vertebrate (Dataset 2). Difference is not statistically significant (p-value = 0.25)

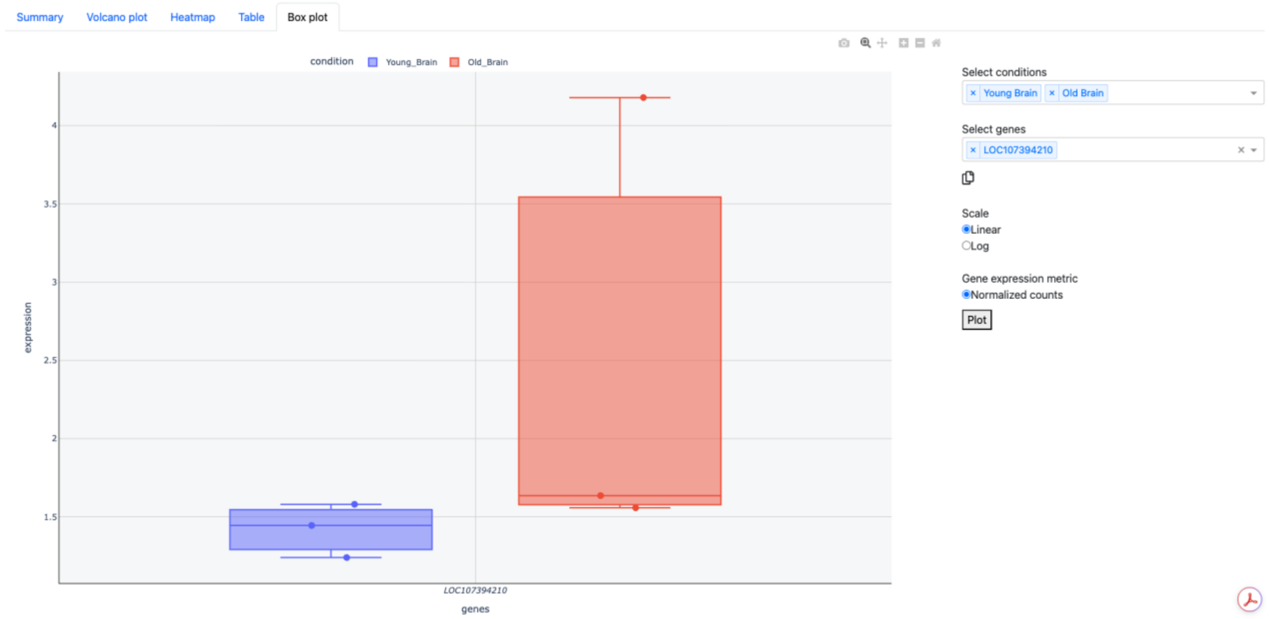

**Figure S13.** UMAP plot for single nucleus RNA-seq dataset from brain in study titled A multi-omic atlas in the African turquoise killifish reveals increased glucocorticoid signaling as a hallmark of brain aging. The points are colored by cell type.

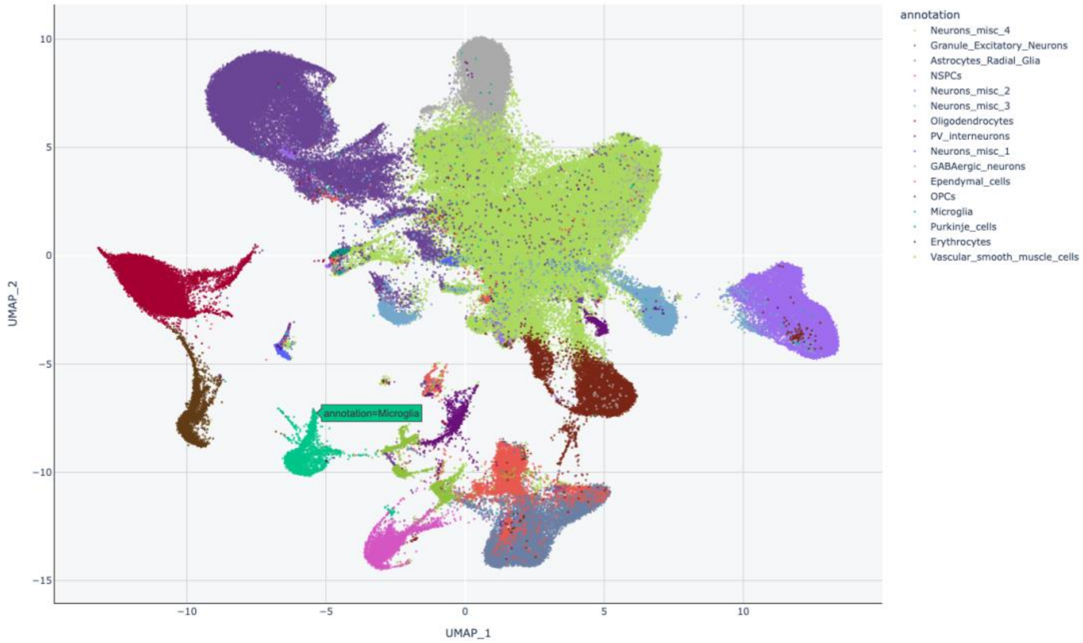


**Figure S14.** UMAP plot for single nucleus RNA-seq dataset from brain in study titled A multi-omic atlas in the African turquoise killifish reveals increased glucocorticoid signaling as a hallmark of brain aging. The points are colored by expression for *LOC107394210*.

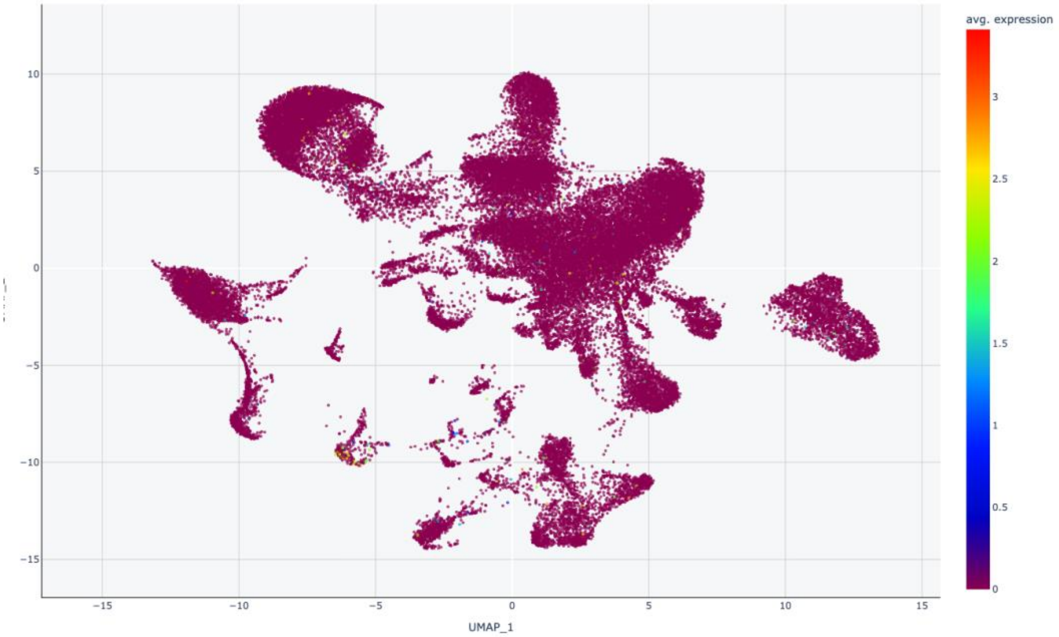

**Figure S15**. Bubble plot for single nucleus RNA-seq dataset from brain in study titled A multi-omic atlas in the African turquoise killifish reveals increased glucocorticoid signaling as a hallmark of brain aging. Bubble size represents the fraction of cells with non-zero expression of *LOC107394210*, while color indicates the average expression level among cells with non-zero expression.

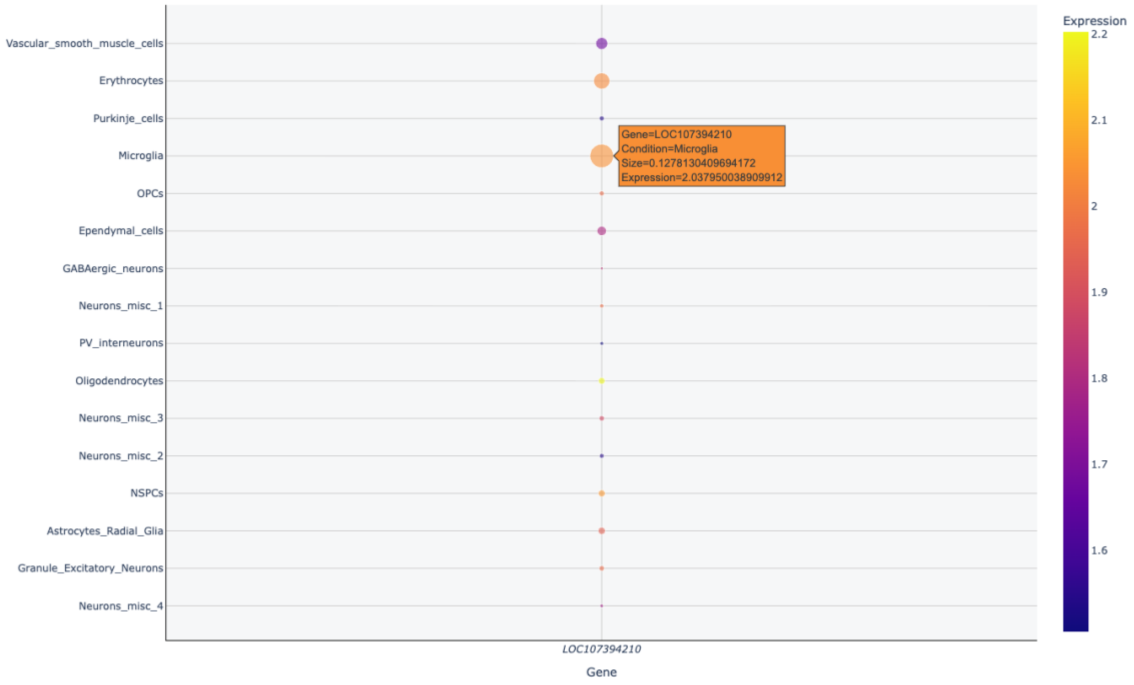
